# Single Cell RNAseq to identify subpopulations of glial progenitors in iPSC-derived oligodendroglial lineage cultures

**DOI:** 10.1101/2024.12.18.629204

**Authors:** Gabriela J Blaszczyk, Valerio E C Piscopo, Taylor M Goldsmith, Alexandra Chapleau, Julien Sirois, Geneviève Bernard, Jack P Antel, Thomas M Durcan

## Abstract

Cellular heterogeneity is a common issue in differentiation protocols of oligodendrocytes (OLs) from human induced pluripotent stem cells. Our previous work described a novel method to generate OLs and highlighted the presence of glial progenitors. Here, we unravel the glial heterogeneity and characterize the response of isolated subpopulations to differentiation. This study provides a novel tool for studying the dynamics of glial development in *vitro* and on a transcriptomic level.

## Main Text

During normal developmental processes, oligodendrocytes (OLs), the myelinating cells of the central nervous system (CNS), differentiate from their precursor cells (OPCs). During development, there are three waves of OPC differentiation, allowing for rapid population of the developing CNS, although not all OPCs will survive until adulthood of the organism (1). Since their discovery (2), many terms have been utilized to describe OPCs including “NG2” cells, “polydendrocytes” and “O2A” cells (1). OPCs remain resident in the CNS once the brain completes development (1, 3, 4), acting as a reservoir to replace lost OLs in case of injury (4) and to serve a number of homeostatic functions (5, 6). Thus, OPCs serve as an important therapeutic target in demyelinating diseases and regenerative medicine.

Early in the glial lineage, tri-potent glial-restricted progenitors (GRPs) give rise to either type-1 astrocytes directly, astrocyte-restricted progenitors (ARP), or bipotent O2A cells. These O2A cells can further differentiate into type-2 astrocytes or OL-fated OPCs (2, 7, 8, 9). The progression from GRPs to specific differentiated cell types requires tightly regulated developmental steps to ensure a balanced production of astroglial and oligodendroglial populations in the CNS. Early studies on GRP and O2A-cells have suggested that certain *in vitro* conditions can bias their differentiation towards either astrocytic or oligodendroglial lineages (2, 10).

The growing interest in OL lineage cells and their roles in disease pathology underscores the need for reliable and reproducible research models. While murine models have yielded valuable insights into OL biology, significant species differences in OL development, myelination and response to injury present challenges in translating findings to humans (11, 12). These limitations are further emphasized by the absence of myelin pathology in many rodent models carrying disease-causing mutations in genes associated with white matter disorders (13). Studying development in a human *in vitro* model allows us to improve our understanding of unique occurrences that may not be estimated in other organisms.

Access to primary human OL-lineage cells is typically limited to pathological samples from patients with late-stage disease or healthy cells derived from human fetal tissue or surgery for epilepsy, which has become increasingly difficult to obtain due to growing restrictions on its use in research. Thus, human induced pluripotent stem cell (iPSC) models are of increasing importance towards advancing the basic understanding of OPC biology and myelination, and for improving the translatability of future therapeutics targeting these cells and their remyelination potential. However, current efforts are limited by the difficulties in obtaining pure OL cultures from human iPSCs. Astrocytic cells, which spontaneously arise from shared OL/astrocyte progenitors, frequently dominate these cultures, impacting the final yield of OLs. To address this, our group recently published a method to generate OL lineage cells from iPSCs more efficiently, (14). Despite the increased yield of mature OLs, early progenitor heterogeneity still remained an issue.

In the present study, we analyzed the pool of progenitors produced at discrete steps of our OL generation protocol. We identified 3 populations (GRPs, ARPs and O2A) and isolated them using a previously tested in-house fluorescence- activated cell sorting (FACS) pipeline and assessed the differentiation potential of each population to confirm the contribution of each subpopulation to the final proportion of cell types. With single-cell RNA sequencing (scRNAseq), we identified three distinct developmental lineages and the distinct molecular mechanisms contributing to each. This study aims to shed light on human developmental dynamics in an *in vitro* model. This information will serve to set a benchmark for the study of complex glial cell dynamics in healthy and disease contexts thereby allowing for the fine-tuning of further works exploring the development and remyelination capacity of these cells.

With this in mind, we generated OL-lineage cells using our previously published protocol (14) (Figure 1A). To study the contribution of various glial precursor populations to our overall cultures, we sequenced our cultures at different timepoints throughout the protocol (Day 75, Day 85 and Day 95) (Figure 1A, 1C). These timepoints comprise the majority of glial progenitors (day 75), committed OPCs (Day 85) and pre-OLs/OLs (Day 95) (Figure 1A). Following unsupervised clustering, we identified populations of GRPs, O2A and ARPs (Figures 1B, 1C) as well as various sub-populations of OPCs and astrocytes using canonical markers (Figures 1B, 1C). Next, we asked which glial clusters responded to our differentiation media, namely at our selected timepoints (Day 75, Day 85 and Day 95) where media composition is changed (see materials and methods). Interestingly, at each of the selected timepoints in the protocol, we observed stable proportions of both proliferative and resting OPC clusters, as well as the O2A and GRP populations (Figure 1D). Clusters with the largest changes in response to the media included “Astro_3”, “Astro_1”, “ARP”, “pre OL” and “late OPC” (Figure 1D). These observations further solidify our group’s previous findings (14), where OL differentiation was observed to coincide with the generation of astrocytes and retention of glial progenitors.

**Figure 1.**
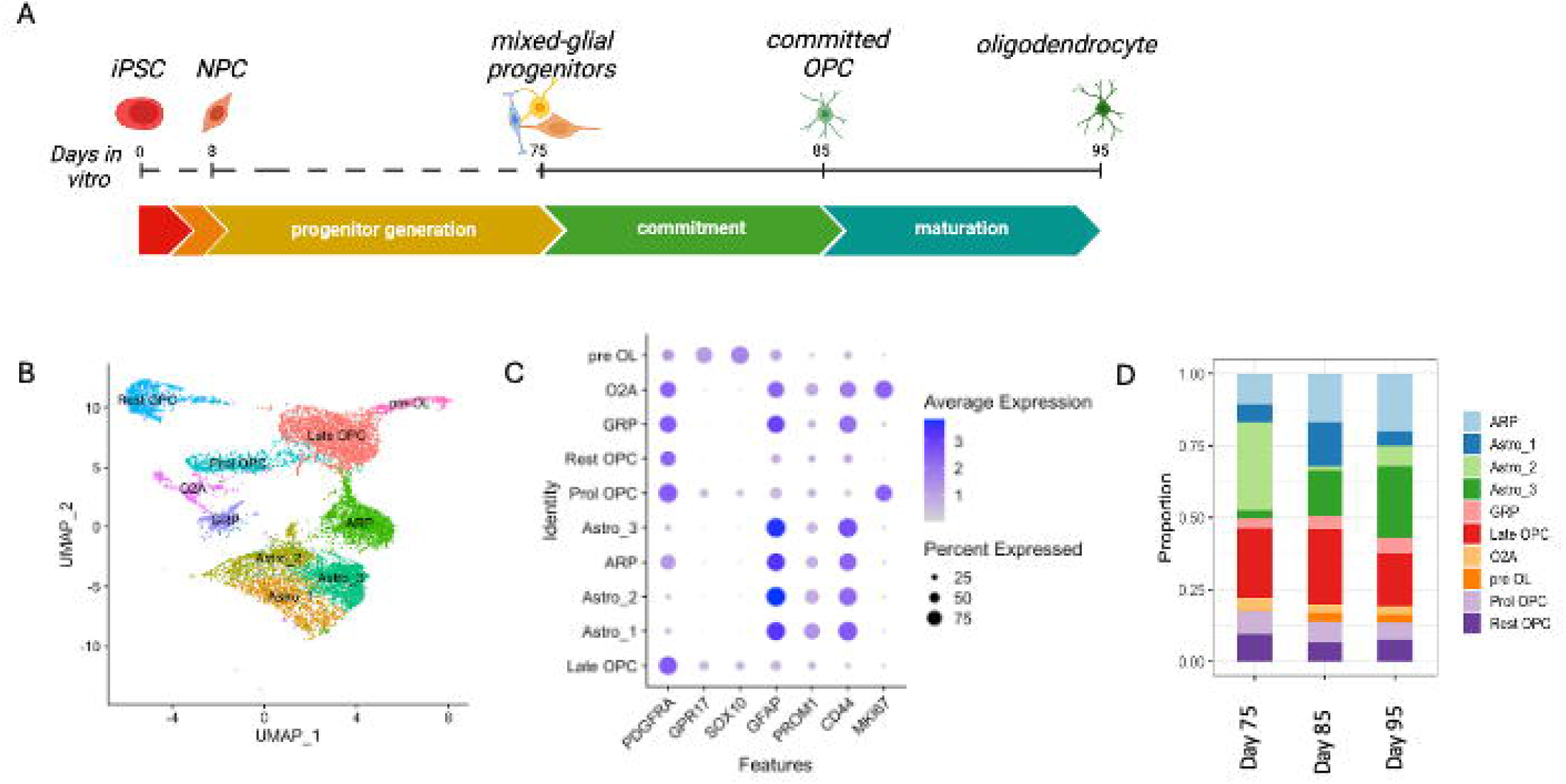
Generation of OL-lineage cells by growth factors results in the retention of glial precursor populations. **(A)** Schematic of the OL-lineage generation protocol from human iPSCs. **(B)** Dimplot following unsupervised clustering of two iPSC lines at Day 75, Day 85 and Day 95 of the protocol following scRNAseq, all shown. (**C)** Dot plot representation the average expression of key identifying genes used to label clusters. **(D)** Bar plot representing cluster proportions at selected stages of the differentiation protocol.

We further assessed the lineage potential of identified glial progenitor populations by employing FACS, isolating them using a combination of antibodies previously described by our lab (14). We utilized combinations of the following: A2B5+PDGFR⍰+ for OPCs, A2B5+PDGFR⍰- for GRP and CD44+PDGFR⍰- for ARP, confirmed in our scRNAseq dataset (Figure 1C). We next cultured the populations in OL maturation media for 21 days followed by immunofluorescence staining to confirm the identity of the glial subpopulations. Specifically, most GRPs expressed astrocytic markers S100β and GFAP (Figures 2 A-B) with a small subpopulation displaying low levels of OPC/OL markers PDGFR⍰ and O4 (Figures 2 C-D). ARPs gave rise almost exclusively to S100β and GFAP-positive cells and displayed high levels of the nuclear factor 1A (NF1A, Figure 2C), associated with astrocytic lineage fate (15). In turn, O2A/OPC cells predominantly differentiated into OLIG2, PDGFR⍰ and O4-positive cells (Figures 2 C-D) with some sparse cells expressing the myelin marker MBP, which was completely absent in the other populations (Figure 2D). Thus, while both GRPs and O2A cells could differentiate to OPCs, only OPCs arising from O2A cells had the capacity to differentiate into mature OLs.

**Figure 2.**
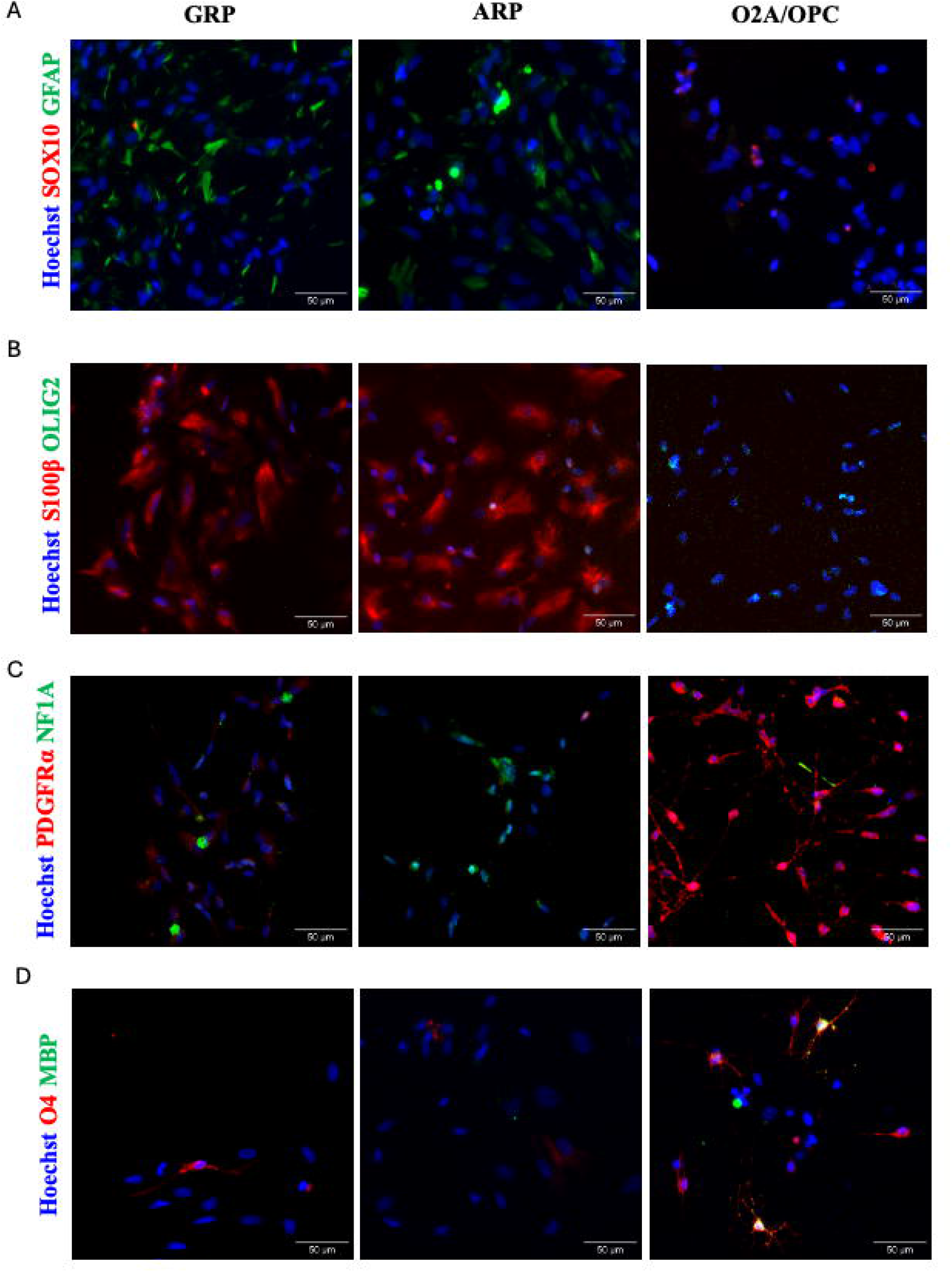
Isolated glial precursor population varied response to differentiation factors. Immunofluoresence staining to characterize sorted glial precursor sub-populations after differentiation. **(A)** SOX10 in red, GFAP in green. **(B)** S100β in red, OLIG2 in green. **(C)** PDGFR⍰ in red, NF1A in green. **(D)** O4 in red, MBP in green.

To explore the cell fate decisions of our glial precursor subpopulations, we analyzed their developmental trajectories in silico (16). Following guided analysis starting at the GRP cluster, we identified three separate lineages, lineage 1 (terminating at astro_3 in our dataset), lineage 2 (rest OPC) and lineage 3 (pre OL) (Figure 3A). Lineages 2 and 3 have been identified as essential OPC subpopulations in the rodent brain (17), which are retained throughout postnatal life. To understand the molecular drivers underlying the divergence of lineages 2 and 3, we evaluated the top differentially expressed genes (Figures 3B-C). The genes were manually curated based on the output of tradeSeq analysis (18), for example *SGCD, RNF144A, DNM3 and PKP4* contributing to lineage 3 (Figure 3C). Our data shows that although OPCs are generated with our differentiation protocol, only a proportion of these are destined to become mature OLs while the other are retained as an OPC reservoir containing both resting and proliferative cells.

**Figure 3.**
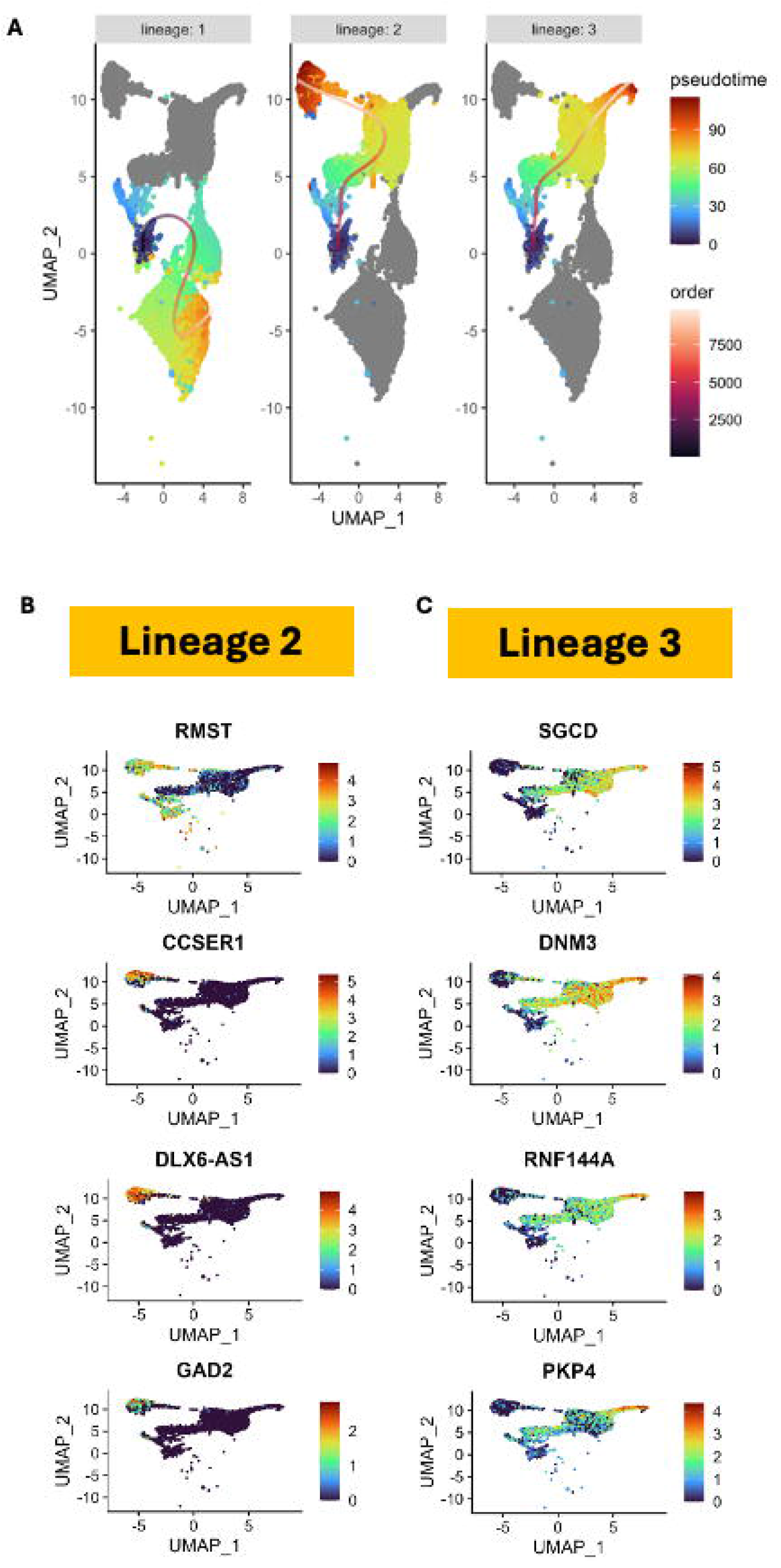
Only one precursor population is destined to give rise to mature OLs. **(A)** Dim plots representing pseudotime lineages as generated by Slingshot originating from the “GRP” cluster. Lineages end at the “Astro_3”, “rest OPC”, and “pre OL” clusters. **(B-C)** The Feature Plots represent the top most significant genes which contribute to lineage 2 **(B)** and lineage 3 **(C)** following subset for OL-lineage cells and patternTest.

The differentiation of OL-lineage cells from glial precursor populations is a dynamic and multifaceted process. The sequencing of cultures at various timepoints (Day 75, 85, and 95) revealed stable populations of OPCs, O2A precursors, and GRPs, suggesting that these progenitors maintain their identities even under varying conditions. The biological meaning of the persistence of these precursors remains to be established and would require further study. Following pseudotime analysis, we observed the retention a cluster of OPCs capable of generating mature OLs as well as a reservoir of resting OPCs. Resting, or homeostatic OPCs (17) have a broad range of functions which are important for CNS development and maintenance. Their roles include angiogenesis (5) and mediation of inflammation within the CNS (6). Additionally, the differential gene expression analysis at the bifurcation point of the two identified lineages offered insight into the molecular mechanisms that drive lineage commitment (Figure 3). The presence of a resting OPCs reservoir can provide a novel target for further works addressing remyelination and human developmental processes.

These results suggest that while multiple glial progenitor populations are present, only a specific subset ultimately gives rise to mature OLs, with the remaining cells retained or generated astrocytes leading to heterogeneity in the dish. Here we provide a workflow for the isolation and interrogation of early OL-lineage cells. Our flow cytometry panel to subdivide glial progenitors provides an avenue for the generation of a purer OL culture by the isolation of O2A cells (Figure 2). Further, by elucidating which transcripts are prominent for fate switching in the OL-lineage (Figure 3), we can then find compounds to aid in generation of late-stage OLs. This work provides a deeper understanding of how glial precursor populations contribute to OL development in human cells, and we provide a novel model for studying myelination, remyelination, and glial cell plasticity in neurological diseases.

## Methods

### Cell Lines

The lines used in this study include 3450 (19) and 81280 (obtained from Genome Quebec). The complete profiles of the iPSCs have been published previously (14). The use of iPSCs and stem cells in this research was approved by the McGill University Health Centre Research Ethics Board (DURCAN_IPSC/2019–5374).

### OL-lineage generation from human iPSCs

Glial progenitors were generated from human iPSCs according to our previous publication (14). At day 75 of the protocol, we have a mixed-glial culture consisting of committed glial progenitors, astrocytes and OPCs (Figure 1B). Passaging our mixed-progenitor culture allows for expansion and use for differentiation as needed. For differentiation, a commercially available basal medium is supplemented with growth serum (Astrocyte Medium with Astrocyte Growth Supplements, ScienCell) for 10 days (Day 85), followed by 10ng/mL IGF-1 (Peprotech) and 60ng/mL T3 (Sigma Aldrich) for the following 11 days to reach a majority of OL-lineage cells at a total of 21 days (Day 95).

### Single Cell Sequencing

#### Library Preparation, Sequencing, Alignment

Cells generated from both 3450 and 81280, were collected and fixed using the Parse fixation workflow that allows samples to be stored at -80°C until the barcoding and library steps are carried out. Between 100,000 and four million cells may enter the fixation. On the day of library preparation, cells are combinatorically barcoded, with each cell receiving three unique barcodes over three consecutive splitting and pooling steps. Barcoded cells are then split into sublibraries, lysed, reverse transcribed, and barcoded once more in a sub library-specific manner. Sublibraries are sequenced through the McGill Genome Centre Platform using an Illumina NovaSeq 6000. Generated Fastq files are read into the Parse Bioscience analysis pipeline on the high-performance Digital Research Alliance of Canada computing cluster Béluga to assign samples to barcodes, combine sublibraries, and undergo initial quality control. *Analysis (Seurat, Slingshot, tradeSeq)* The Parse pipeline output was read into R v4.3.3 using Seurat v5.1.0 to further quality control and process the data. Slingshot v2.10.0 (16) was used for the pseudotime analysis, and tradeSeq v1.16.0 (18) for evaluation of DEGs associated with lineage divergence.

### Fluorescence Activated Cell Sorting

Live sorting was performed using the aria fusion cell sorter (BD Biosciences). Cells were collected and passed through 40 µM mesh to ensure single cell suspension. Cells were stained with CD44 (Biolegend), PDGFR⍰ (BD Optibuild) and A2B5 (Miltenyi biotec) as previously described (14). Cells were placed into culture post-FACS for differentiation and analysis by immunofluorescence imaging.

### Immunofluorescence Staining and Imaging

Cells were stained as previously described (14). Briefly, cells were stained with Anti-O4 (R&D), SOX10 (R&D), PDGFR⍰ (Cell Signaling), OLIG2 (Millipore), NF1A (Sigma), GFAP (Dako) and S100β (Sigma) to characterize cells resulting from differentiation post-FACS. Images were acquired with a Zeiss Axio Observer Z1 Inverted Microscope using 20x magnification objective (N.A 0.8) and a ZEISS Axiocam 506 mono camera.

## Acknowledgements

We thank lab members Irina Shlaifer, Wolfgang Reintsch, Andrea Krahn, and Genevieve Dorval for technical or administrative assistance. Thank you to Carol Chen, Michael Nicouleau, and Aeshah Alluli for assistance in processing of samples and library preparation for scRNAseq. Thank you to Elia Afanasiev for assistance with the Slingshot/TradeSeq workflow. The flow cytometry work and cell sorting were performed at The Neuro’s Early Drug Discovery Unit’s Flow Cytometry Core Facility. We acknowledge the McGill microscopy platform for their support. Sequencing was done at the McGill Genome Centre’s McGill Applied Genomics Innovation Core (MAGIC). GB has received the Clinical Research Scholar Junior 1 Award from the Fonds de Recherche du Québec-Santé (FRQS) (2012-2016), the New Investigator Salary Award from the Canadian Institutes of Health Research (CIHR) (2017- 2022), the Clinical Research Scholar Senior Award from the FRQS (2022-2025), and the Chercheur de Mérite Award from the FRQS (2025-2029). TMD received funding to support this project through the Canada First Research Excellence Fund, awarded through the Healthy Brains, Healthy Lives initiative at McGill University and an ALS Canada/Brain Canada Discovery grant.

## Author Contributions

GJB and VECP contributed substantially to the conception and design of the study. GJB, VECP, AC and TMG drafted a significant portion of the manuscript or figures. GJB, VECP, AC, TMG and JS contributed to acquisition and analysis of data. JPA, GB and TMD contributed to critical manuscript revision and oversaw the project. All authors read and approved the final version of the manuscript.

## Data Availability

The datasets generated and/or analyzed during the current study will be available in the Gene Expression Omnibus (GEO) upon acceptance/publication of the manuscript.

## Code Availability

The underlying code for this study will be available on Github and can be accessed following acceptance/publication of the manuscript.

## References

1. Bergles DE, Richardson WD. Oligodendrocyte Development and Plasticity. Cold Spring Harb Perspect Biol. 2015;8(2):a020453.

2. Raff MC, Abney ER, Cohen J, Lindsay R, Noble M. Two types of astrocytes in cultures of developing rat white matter: differences in morphology, surface gangliosides, and growth characteristics. J Neurosci. 1983;3(6):1289–300.

3. Reynolds R, Hardy R. Oligodendroglial progenitors labeled with the O4 antibody persist in the adult rat cerebral cortex in vivo. J Neurosci Res. 1997;47(5):455–70.

4. Crawford AH, Stockley JH, Tripathi RB, Richardson WD, Franklin RJ. Oligodendrocyte progenitors: adult stem cells of the central nervous system? Exp Neurol. 2014;260:50–5.

5. Yuen TJ, Silbereis JC, Griveau A, Chang SM, Daneman R, Fancy SPJ, et al. Oligodendrocyte-encoded HIF function couples postnatal myelination and white matter angiogenesis. Cell. 2014;158(2):383–96.

6. Moyon S, Dubessy AL, Aigrot MS, Trotter M, Huang JK, Dauphinot L, et al. Demyelination causes adult CNS progenitors to revert to an immature state and express immune cues that support their migration. J Neurosci. 2015;35(1):4–20.

7. Rao MS, Mayer-Proschel M. Glial-restricted precursors are derived from multipotent neuroepithelial stem cells. Dev Biol. 1997;188(1):48–63.

8. Gregori N, Pröschel C, Noble M, Mayer-Pröschel M. The tripotential glial-restricted precursor (GRP) cell and glial development in the spinal cord: generation of bipotential oligodendrocyte-type-2 astrocyte progenitor cells and dorsal-ventral differences in GRP cell function. J Neurosci. 2002;22(1):248–56.

9. Mi H, Barres BA. Purification and characterization of astrocyte precursor cells in the developing rat optic nerve. J Neurosci. 1999;19(3):1049–61.

10. Raff MC, Abney ER, Miller RH. Two glial cell lineages diverge prenatally in rat optic nerve. Dev Biol. 1984;106(1):53–60.

11. Bradl M, Lassmann H. Oligodendrocytes: biology and pathology. Acta Neuropathol. 2010;119(1):37–53.

12. Jakovcevski I, Filipovic R, Mo Z, Rakic S, Zecevic N. Oligodendrocyte development and the onset of myelination in the human fetal brain. Front Neuroanat. 2009;3:5.

13. Choquet K, Yang S, Moir RD, Forget D, Larivière R, Bouchard A, et al. Absence of neurological abnormalities in mice homozygous for the Polr3a G672E hypomyelinating leukodystrophy mutation. Mol Brain. 2017;10(1):13.

14. Piscopo VEC, Chapleau A, Blaszczyk GJ, Sirois J, You Z, Soubannier V, et al. The use of a SOX10 reporter toward ameliorating oligodendrocyte lineage differentiation from human induced pluripotent stem cells. Glia. 2024;72(6):1165–82.

15. Deneen B, Ho R, Lukaszewicz A, Hochstim CJ, Gronostajski RM, Anderson DJ. The Transcription Factor NFIA Controls the Onset of Gliogenesis in the Developing Spinal Cord. Neuron. 2006;52(6):953–68.

16. Street K, Risso D, Fletcher RB, Das D, Ngai J, Yosef N, et al. Slingshot: cell lineage and pseudotime inference for single-cell transcriptomics. BMC Genomics. 2018;19(1):477.

17. Dennis DJ, Wang BS, Karamboulas K, Kaplan DR, Miller FD. Single-cell approaches define two groups of mammalian oligodendrocyte precursor cells and their evolution over developmental time. Stem Cell Reports. 2024;19(5):654–72.

18. Van den Berge K, Roux de Bézieux H, Street K, Saelens W, Cannoodt R, Saeys Y, et al. Trajectory-based differential expression analysis for single-cell sequencing data. Nature Communications. 2020;11(1):1201.

19. Chen CX, Abdian N, Maussion G, Thomas RA, Demirova I, Cai E, et al. A Multistep Workflow to Evaluate Newly Generated iPSCs and Their Ability to Generate Different Cell Types. Methods Protoc. 2021;4(3).

